# TreeSearch: morphological phylogenetic analysis in R

**DOI:** 10.1101/2021.11.08.467735

**Authors:** Martin R. Smith

## Abstract

TreeSearch is an R package for phylogenetic analysis, optimized for morphological datasets. Tree search may be conducted using equal or implied step weights with an explicit (albeit inexact) allowance for inapplicable character entries, avoiding some of the pitfalls inherent in standard parsimony methods. Profile parsimony and user-specified optimality criteria are supported.

A graphical interface, which requires no familiarity with R, is designed to help a user to improve the quality of datasets through critical review of underpinning character codings; and to obtain additional information from results by identifying and summarizing clusters of similar trees, mapping the distribution of trees, and removing ‘rogue’ taxa that obscure underlying relationships.

Taken together, the package aims to support methodological rigour at each step of data collection, analysis, and the exploration of phylogenetic results.

---

Even in the phylogenomic era, morphological data make an important contribution to phylogenetic questions. Phenotypic data improve the accuracy and resolution of phylogenetic reconstruction even when outnumbered by molecular characters, and are the only way to incorporate the unique perspective on historical events that fossil taxa provide (Wiens 2004; Wortley and Scotland 2006; Koch and Parry 2020; Asher and Smith 2022).

One challenge with morphological analysis is the treatment of inapplicable character states: for example, ‘tail colour’ cannot logically be ascribed either of the states ‘red’ or ‘blue’ in a taxon that lacks a tail (Maddison 1993). This situation can profoundly mislead phylogenetic analysis, and is not handled appropriately by any standard Markov model or parsimony method.

Solutions to this issue have recently been proposed (De Laet 2005; Brazeau et al. 2019; Tarasov 2019; Goloboff et al. 2021; Hopkins and St. John 2021). Where a single ‘principal’ character (e.g. ‘tail’) exhibits *n*’contingent’ characters (e.g. ‘tail colour,’ ‘tail covering’), ‘exact’ solutions (Tarasov 2019; Goloboff et al. 2021) require the construction of multi-state hierarchies containing *O* (2^*n*^) entries, meaning that analysis is only computationally tractable for simple hierarchies with few contingent characters. Moreover, these approaches cannot accommodate characters that are contingent on more than one principal character: for example, characters describing appendages on a differentiated head may be contingent on the presence of the two characters ‘appendages’ and ‘differentiated head.’

Such situations can be approached using the flexible but approximate parsimony approach proposed by Brazeau et al. (2019). TreeSearch scores trees using the “Morphy” C implementation of this algorithm (Brazeau et al. 2017). Morphy implements tree search under equal step weights. TreeSearch additionally implements implied step weighting (Goloboff 1993), a method which consistently finds more accurate and precise trees than equal weights parsimony (Goloboff et al. 2008, 2018a; Smith 2019a).

There has been lively discussion as to whether, with the rise of probabilistic approaches, parsimony remains a useful tool for morphological phylogenetics (e.g. O’Reilly et al. 2016; Puttick et al. 2017; Brown et al. 2017; Sansom et al. 2018; Goloboff et al. 2018b).

Notwithstanding scenarios that go beyond the limits of parsimony, such as the simultaneous incorporation of stratigraphic data and other prior knowledge (e.g. Guenser et al. 2021), neither parsimony nor probabilistic methods consistently recover ‘better’ trees when gains in accuracy are balanced against losses in precision (Smith 2019a). Even if probabilistic methods may eventually be improved through the creation of more sophisticated models that better reflect the nature of morphological data (Goloboff et al. 2018a; Tarasov 2019), parsimony analysis remains a useful tool – not only because treatments of inapplicable character states are presently available, but also because it facilitates a deeper understanding of the underpinning data by emphasizing the reciprocal relationship between a tree and the synapomorphies that it implies.

Whatever method is used to find phylogenetic trees, a single consensus tree may fail to convey all the signal in a set of phylogenetic results (Wilkinson 1994, 1996, 2003). A set of optimal trees can be better interpreted by examining consensus trees generated from clusters of similar trees (Stockham et al. 2002); by exploring tree space (Wright and Lloyd 2020; Smith 2022a) and by automatically identifying, annotating and removing ‘wildcard’ taxa (Smith 2022b) whose ‘rogue’ behaviour may reflect underlying character conflict or ambiguity (Kearney 2002). These methods are not always easy to integrate into phylogenetic workflows, so are not routinely included in empirical studies.

TreeSearch provides functions that allow researchers to engage with the three main aspects of morphological phylogenetic analysis: dataset construction and validation; phylogenetic search (including with inapplicable data); and the interrogation of optimal tree sets. These functions can be accessed through the R command-line, as documented within the package and at ms609.github.io/TreeSearch/, or via an integrated graphical user interface (GUI), with options to save outputs in graphical formats or as Nexus or Newick files for further analysis.

## Implementation

### Dataset review

Ultimately, the quality of a dataset plays a central role in determining the reliability of phylogenetic results, with changes to a relatively small number of character codings potentially exhibiting an outsized impact on reconstructed topologies (Goloboff and Sereno 2021).

Nevertheless, dataset quality does not always receive commensurate attention (Simões et al. 2017). One step towards improving the rigour of morphological datasets is to annotate each cell in a dataset with an explicit justification for each taxon’s coding (Sereno 2009), which can be accomplished in Nexus-formatted data files (Maddison et al. 1997) using software such as MorphoBank (O’Leary and Kaufman 2011).

TreeSearch presents such annotations alongside a reconstruction of each character’s states on an optimal tree, with inapplicable states mapped according to the algorithm of Brazeau et al. (2019). Neomorphic (presence/absence) and transformational characters (Sereno 2007) are distinguished by reserving the token `θ` to solely denote the absence of a neomorphic character, with tokens `1` … `n` used to denote the *n*states of a transformational character (Brazeau et al. 2019).

This visualization of reconstructed character transitions can help to identify cases where the formulation of characters has unintended consequences (Wilkinson 1995; Brazeau 2011); where inapplicable states have been inconsistently applied (Brazeau et al. 2019); where taphonomic absence is wrongly coded as biological absence (Donoghue and Purnell 2009); where previous datasets are uncritically recycled (Jenner 2001); or where taxa are coded with more confidence than a critical evaluation of available evidence can truly support. Insofar as the optimal tree and the underlying characters are reciprocally illuminating (Mooi and Gill 2016), successive cycles of phylogenetic analysis and character re-formulation can improve the integrity of morphological datasets, and thus increase their capacity to yield meaningful phylogenetic results (Hennig 1966).

### Tree search

The TreeSearch GUI uses the routine `MaximizeParsimony()` to search for optimal trees using tree bisection and reconnection (TBR) searches and the parsimony ratchet (Nixon 1999). This goes beyond the heuristic tree search implementation in the R package “phangorn” (Schliep 2011) by using compiled C code to rearrange trees, accelerating computation; and in supporting TBR rearrangements, which explore larger neighbourhoods of tree space: TBR evaluates more trees than nearest-neighbour interchanges or subtree pruning and regrafting, leading to additional computational expense that is offset by a decreased likelihood that search will become trapped in a local optimum (Goeffon et al. 2008; Whelan and Money 2010).

By default, search begins from a greedy addition tree generated by function `AdditionTree()`, which queues taxa in a random order, then attaches each taxon in turn to the growing tree at the most parsimonious location. Search may also be started from neighbour-joining trees, or the results of a previous search.

Search commences by conducting TBR rearrangements – a hill-climbing approach that locates a locally optimal tree from which no tree accessible by a single TBR rearrangement has a better score. A TBR iteration breaks a randomly selected edge in the focal tree, and reconnects each possible pair of edges in the resultant sub-trees to produce a list of candidate trees. Entries that are inconsistent with user-specified topological constraints are removed; remaining trees are inserted into a queue and scored in a random sequence. If the score of a candidate tree is at least as good as the best yet encountered (within the bounds of an optional tolerance parameter ϵ, which allows the retention of almost-optimal trees in order to improve accuracy (e.g. Smith 2019a)), this tree is used as the starting point for a new TBR iteration. Otherwise, the next tree in the list is considered. TBR search continues until the best score is found a specified number of times; a specified number of TBR break points have been evaluated without any improvement to tree score; or a set amount of time has passed.

When TBR search is complete, iterations of the parsimony ratchet (Nixon 1999) are conducted in order to search areas of tree space that are separated from the best tree yet found by ‘valleys’ that cannot be traversed by TBR rearrangements without passing through trees whose optimality score is below the threshold for acceptance. Each ratchet iteration begins by resampling the original matrix. A round of TBR search is conducted using this resampled matrix, and the tree thus produced is used as a starting point for a new round of TBR search using the original data.

After a specified number of ratchet iterations, an optional final round of TBR search allows a denser sampling of optimal trees from the final region of tree space.

During tree search, all trees whose score is within *∈* of the best score are retained, and tagged with the iteration in which they were first identified. This allows the progress of tree search to be visualized in tree space (Fig. 1; after Whidden and Matsen 2015).

**Figure 1.**
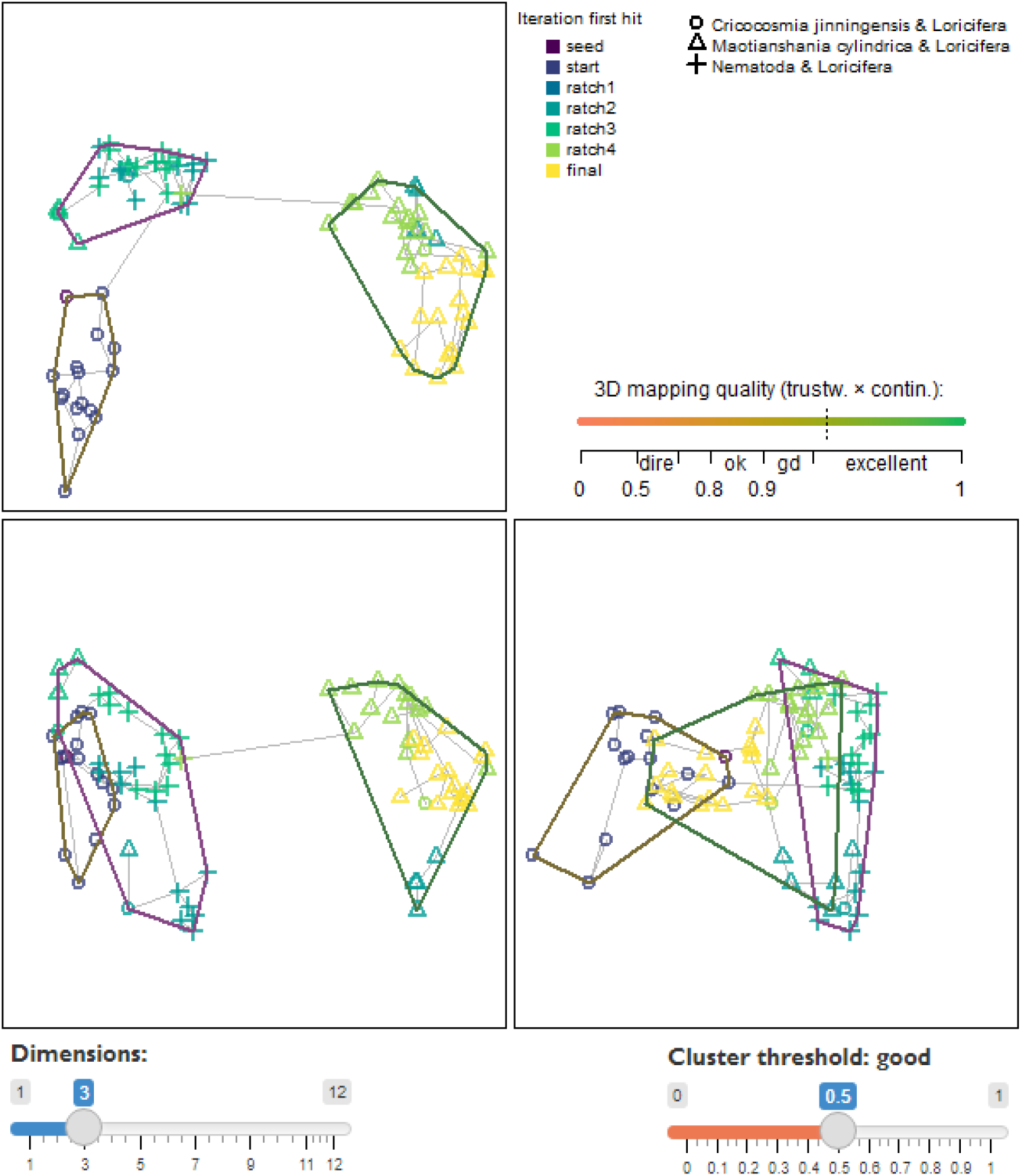
Three-dimensional map visualizing tree search progress. Optimal trees belong to three statistically distinct clusters with good support (silhouette coefficient > 0.5), characterized by different relationships between certain taxa (plotting symbols). Although multiple ratchet iterations have visited each cluster, limited overlap between ratchet iterations suggests that a continuation of tree search may sample novel optimal trees. High trustworthiness and continuity values and a simple minimum spanning tree (grey) indicate that the mapping does not exhibit severe distortion.

More flexible, if less computationally efficient, tree searches can be conducted at the command line using the `TreeSearch()`, `Ratchet()` and `Bootstrap()` commands, which support custom tree optimality criteria (e.g. Hopkins and St. John 2021).

### Tree scoring

Trees may be scored using equal weights, implied weighting (Goloboff 1993), or profile parsimony (Faith and Trueman 2001). The function `TreeLength()` calculates tree score using the “Morphy” phylogenetic library (Brazeau et al. 2017), which implements the Fitch (1971) and Brazeau et al. (2019) algorithms. Morphy returns the equal weights parsimony score of a tree against a given dataset. Implied weights and profile parsimony scores are computed by first making a separate call to Morphy for each character in turn, passed as a single-character dataset; then passing this value to the appropriate weighting formula and summing the total score over all characters.

Implied weighting is an approximate method that treats each additional step in a character as less surprising – and thus requiring less penalty – than the previous step. Each additional step demonstrates that a character is less reliable for phylogenetic inference, and thus more likely to contain additional homoplasy. The score of a tree under implied weighting is 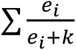, where *e*_*i*_ denotes the number of extra steps observed in character *i*, and is derived by subtracting the minimum score that the character can obtain on any tree from the score observed on the tree in question (Goloboff 1993). The minimum length of a tree is one less than the number of unique applicable tokens that must be present.

Profile parsimony (Faith and Trueman 2001) represents an alternative formulation of how surprising each additional step in a character is (Arias and Miranda-Esquivel 2004): the penalty associated with each additional step in a character is a function of the probability that a character will fit at least as well as is observed on a uniformly selected tree. On this view, an additional step is less surprising if observed in a character where there are more opportunities to observe homoplasy, whether because a character contains fewer ambiguous codings (a motivation for the ‘extended’ implied weighting of Goloboff (2014)) or because states are distributed more evenly in a character, whose higher phylogenetic information content (Thorley et al. 1998) corresponds to a lower proportion of trees in which no additional steps are observed.

TreeSearch calculates the profile parsimony score by computing the logarithm of the number of trees onto which a character can be mapped using *m* steps, using theorem 1 of Carter et al. (1990). As computation for higher numbers of states (Maddison and Slatkin 1991) is more computationally complex, the present implementation is restricted to characters that contain two applicable tokens, and uses the Fitch (1971) algorithm.

### Visualization

The distribution of optimal trees, however obtained, can be visualized interactively through mappings of tree space (Hillis et al. 2005; Smith 2022a) using the TreeSearch GUI.

The GUI supports the use of information theoretic distances (Smith 2020a); the quartet distance (Estabrook et al. 1985); or the Robinson–Foulds distance (Robinson and Foulds 1981) to construct tree spaces, which are mapped into 2–12 dimensions using principal coordinates analysis (Gower 1966). The degree to which a mapping faithfully depicts original tree-to-tree distances is measured using the product of the trustworthiness and continuity metrics (Venna and Kaski 2001; Kaski et al. 2003; Smith 2022a), a composite score denoting the degree to which points that are nearby when mapped are truly close neighbours (trustworthiness), and the degree to which nearby points remain nearby when mapped (continuity). A visualization of stress is provided by plotting the minimum spanning tree (Gower and Ross 1969); contortions in this tree indicate that mapping has distorted original distances (Smith 2022a).

To relate the geometry of tree space to the underlying trees, each point in tree space may be annotated according to the optimality score of its corresponding tree under a selected step weighting scheme; by the relationships between chosen taxa that are inferred by that tree; and by the search iteration in which the tree was first found by tree search (Fig. 1).

The latter feature can be used to evaluate whether a continuation of tree search is likely to yield more optimal trees. For example, if the trees retained are first found only in later rounds of tree search, this recent improvement in tree score suggests that a global optimum may not yet have been reached. Alternatively, if each individual ratchet iteration samples a separate region of tree space, it is likely that the landscape of optimal trees contains isolated ‘islands’ (Bastert et al. 2002), some of which may remain to be found. Continuing tree search until additional ratchet iterations no longer locate new clusters of trees will reduce the chance that optimal regions of tree space remain unvisited.

As the identification of clusters from mappings of tree space can be misleading (Smith 2022a), TreeSearch identifies clusters of trees from tree-to-tree distances using K-means clustering, partitioning around medoids and hierarchical clustering with minimax linkage (Hartigan and Wong 1979; Murtagh 1983; Bien and Tibshirani 2011; Maechler et al. 2019). Clusterings are evaluated using the silhouette coefficient, a measure of the extent of overlap between clusters (Kaufman and Rousseeuw 1990). The clustering with the highest silhouette coefficient is depicted if the silhouette coefficient exceeds a user-specified threshold; the interpretation of the chosen threshold according to Kaufman and Rousseeuw (1990) is displayed to the user. Plotting a separate consensus tree for each cluster often reveals phylogenetic information that is concealed by polytomies in the single ‘plenary’ consensus of all optimal trees (Stockham et al. 2002).

Plenary consensus trees can also lack resolution because of wildcard or ‘rogue’ taxa, in which conflict or ambiguity in their character codings leads to an unsettled phylogenetic position (Wilkinson 1994, 2003; Kearney 2002). TreeSearch detects rogue taxa using a heuristic approach (Smith 2022b) that seeks to maximize the phylogenetic information content (*sensu* Thorley et al. 1998) of a consensus tree created after removing rogue taxa from input trees. The position of an excluded taxon is portrayed by shading each edge or node of the consensus according to the number of times the specified taxon occurs at that position on an underlying tree (Fig. 2; after Klopfstein and Spasojevic 2019), equivalent to the ‘branch attachment frequency’ of “Phyultity” (Smith and Dunn 2008).

**Figure 2.**
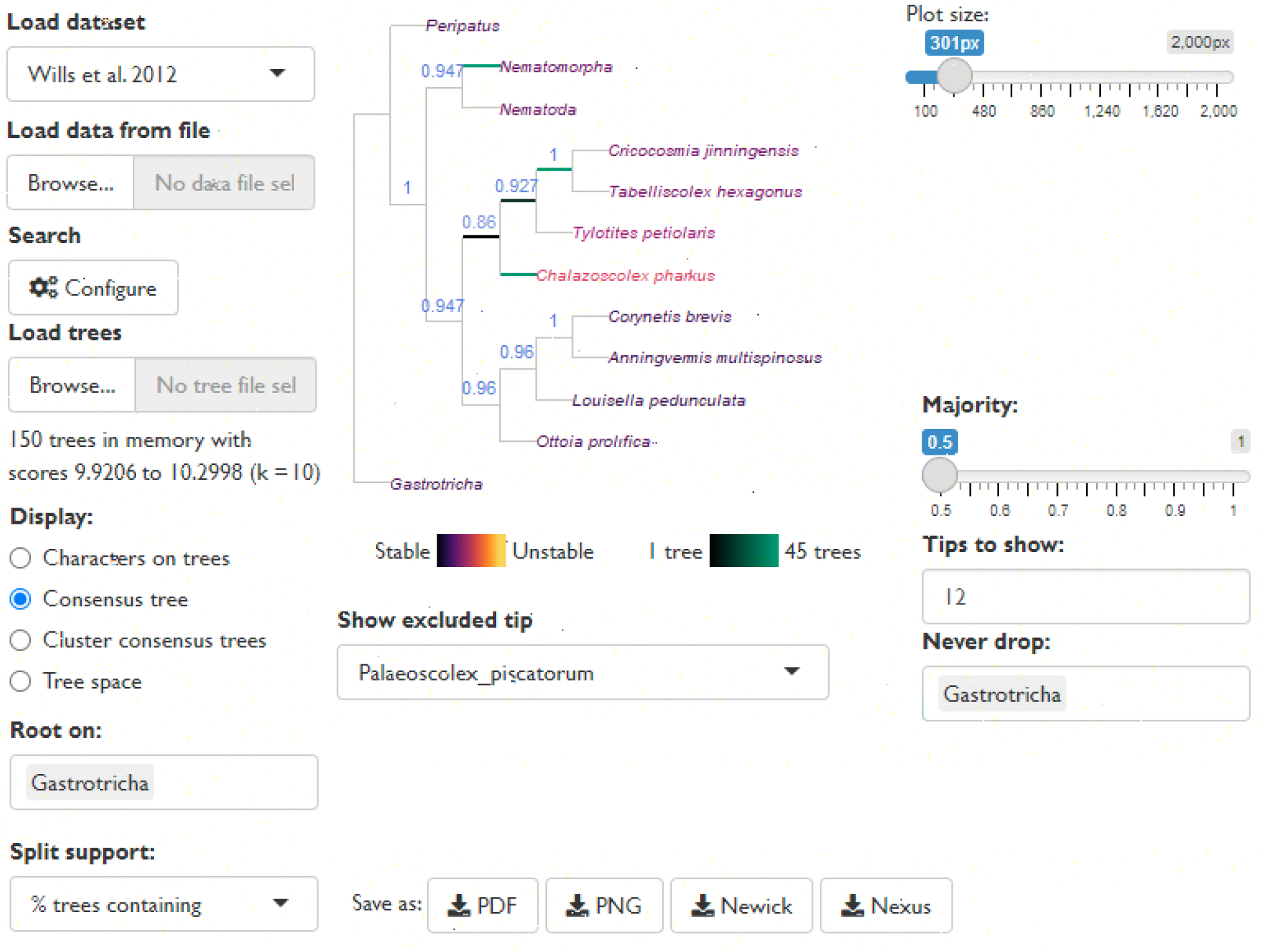
Reduced consensus of 150 cladograms generated by analysis of data from Wills et al. (2012) under different parsimony methods by Brazeau et al. (2019), as displayed in the TreeSearch graphical user interface. Removal of taxa reveals strong support for relationships that would otherwise be masked by rogues such as *Palaeoscolex*, whose position in optimal trees is marked by the highlighted edges.

Identifying taxa with an unstable position, and splits with low support, can help an investigator to critically re-examine character codings; to this end, each edge of the resulting consensus can be annotated with the frequency of the split amongst the tree set, or with a concordance factor (Minh et al. 2020) denoting the strength of support from the underlying dataset.

## Availability

TreeSearch can be installed through the Comprehensive R Archive Network (CRAN) using `install.packages(“TreeSearch”)`; the graphical user interface is launched with the command `TreeSearch::EasyTrees()`. It has been tested on all widely available operating systems and requires only R packages available from the CRAN repository. Source code is available at https://github.com/ms609/TreeSearch/, and is permanently archived at Zenodo (https://dx.doi.org/10.5281/zenodo.1042590). Documentation is online at https://ms609.github.io/TreeSearch/.

~~~
# Note: A temporary bug in the underlying package ‘ape’ v5.6 causes
# issues reading trees from Nexus files.
# A patched version should be installed before installing TreeSearch:
install.packages(“remotes”)remotes::install_github(“ms609/ape@patch-3”)
install.packages(“TreeSearch”)
# The GUI can then be launched with:
library(“TreeSearch”)
EasyTrees()
~~~

## Acknowledgements

I thank Alavya Dhungana and Joe Moysiuk for feedback on preliminary versions of the software, and Martin Brazeau for comments on a draft of the manuscript. Functionality in TreeSearch employs the underlying R (R Core Team 2021) packages ape (Paradis and Schliep 2019), phangorn (Schliep 2011), Quartet (Sand et al. 2014; Smith 2019b), Rogue (Smith 2022b), shiny (Chang et al. 2021), shinyjs (Attali 2020), TreeDist (Smith 2020b), and TreeTools (Smith 2019c).

